# Comparative transcriptome analysis reveals the intensive early-stage responses of host cells to SARS-CoV-2 infection

**DOI:** 10.1101/2020.04.30.071274

**Authors:** Jiya Sun, Fei Ye, Aiping Wu, Ren Yang, Mei Pan, Jie Sheng, Wenjie Zhu, Longfei Mao, Ming Wang, Baoying Huang, Wenjie Tan, Taijiao Jiang

**Affiliations:** Center for Systems Medicine, Institute of Basic Medical Sciences, Chinese Academy of Medical Sciences & Peking Union Medical College, 100005 Beijing, China; Key Laboratory of Medical Virology, National Health and Family Planning Commission, National Institute for Viral Disease Control and Prevention, China CDC, Beijing 102206, China; Suzhou Institute of Systems Medicine, Suzhou, 215123 Jiangsu, China; Department of Otolaryngology, Head and Neck Surgery, Beijing TongRen Hospital, Capital Medical University, Beijing 100730, China

## Abstract

Severe acute respiratory syndrome coronavirus 2 (SARS-CoV-2) has caused a widespread outbreak of highly pathogenic COVID-19. It is therefore important and timely to characterize interactions between the virus and host cell at the molecular level to understand its disease pathogenesis. To gain insights, we performed high-throughput sequencing that generated time-series data simultaneously for bioinformatics analysis of virus genomes and host transcriptomes implicated in SARS-CoV-2 infection. Our analysis results showed that the rapid growth of the virus was accompanied by an early intensive response of host genes. We also systematically compared the molecular footprints of the host cells in response to SARS-CoV-2, SARS-CoV and MERS-CoV. Upon infection, SARS-CoV-2 induced hundreds of up-regulated host genes hallmarked by a significant cytokine production followed by virus-specific host antiviral responses. While the cytokine and antiviral responses triggered by SARS-CoV and MERS-CoV were only observed during the late stage of infection, the host antiviral responses during the SARS-CoV-2 infection were gradually enhanced lagging behind the production of cytokine. The early rapid host responses were potentially attributed to the high efficiency of SARS-CoV-2 entry into host cells, underscored by evidence of a remarkably up-regulated gene expression of TPRMSS2 soon after infection. Taken together, our findings provide novel molecular insights into the mechanisms underlying the infectivity and pathogenicity of SARS-CoV-2.

## Introduction

Coronavirus disease 2019 (COVID-19) triggered by the severe acute respiratory syndrome coronavirus 2 (SARS-CoV-2) is currently affecting global health. The SARS-CoV-2 is the third highly pathogenic coronavirus following SARS-CoV and MERS-CoV that cause severe accurate respiratory symptoms in humans. Since December 2019, this virus has caused more than 80 thousand COVID-19 cases in China. Nowadays, the number of infections in countries outside China is growing rapidly. The most remarkable feature of the SARS-CoV-2 incidences and epidemiology is its great capacity for human-to-human transmission[1]. Clinically, the majority of COVID-19 patients have mild and moderate symptoms, and the elderly appear to have severe symptoms [2]. Based on the analysis of China data, the COVID-19 case-fatality rate was estimated as around 4.0% (3,341deaths over 82,249 confirmed cases of SARS-CoV-2 infection)[3], lower than those of SARS and MERS[4]. However, due to the large-scale infected population, the SARS-CoV-2 has already caused more than ninety thousand deaths as of April 11^th^ 2020, sowing great social panic around the world.

While recent efforts have been focused on transcriptome analysis of host responses to SARS-CoV-2 infection at a certain time point in certain cell lines[5, 6], the transcriptional dynamics of host responses to the virus infection has remained largely unexplored. Generally, once the virus enters the cell, the host innate immune responses, such as the interferon-mediated antiviral responses and cytokine production, have a pivotal role in suppressing the virus replication, which, if inadequate, might contribute to the viral pathogenesis. This hypothesis has been supported by our previous study, which has shown that the high pathogenicity of avian influenza virus is associated with abnormal coordination between interferon-mediated antiviral responses and cytokine production in host cells [7]. Similar to both SARS-CoV and MERS-CoV, which induce the overactivation of cytokines [8], increased cytokine levels are also observed in patients hospitalized with COVID-19 [9]. Transcriptome analysis of *in vitro* host cells shows that SARS-CoV and MERS-CoV elicit distinct responses to the expression of the host genes [10]. Until now, the time-series gene-expression profiling of the host response to SARS-CoV-2 remains unknown and thus is urgently needed uncovering its pathogenesis.

In this study, we used the SARS-CoV-2 strain isolated from patients[11] to infect *in vitro* Calu-3 cells, and performed RNA sequencing to determine the time-series transcriptome profiling data of the host. We established the host response patterns for SARS-CoV-2 by comprehensive analysis of the transcriptomic profiles from SARS-CoV-2, SARS-CoV and MERS-CoV. These results provide profound new insights into the pathogenesis and progression of the COVID-19 disease caused by SARS-CoV-2, illuminating new strategies for the prevention and control of SARS-CoV-2 transmission and eventually leading to a cure of the COVID-19 disease.

## Materials and Methods

### Cells and virus

Calu-3 human airway epithelial cells(ATCC, HTB-55) were cultured in minimum essential media (MEM, HyClone) supplemented with 10% fetal bovine serum(FBS), 1% MEM NEAA, and 100 U/mL penicillin-streptomycin solution (Gibco, Grand Island, USA) at 37 °C in a humidified atmosphere of 5% (v/v) CO_2_. Vero cells (ATCC, CCL-81) were cultured at 37 °C in Dulbecco’s modified Eagle’s medium (DMEM, Gibco) supplemented with 10% FBS (Gibco) in the atmosphere with 5% CO_2_. SARS-CoV-2 strain BetaCoV/Wuhan/IVDC-HB-01/2019 (C-Tan-HB01, GISAID accession no. EPI_ISL_402119) was isolated from a human patient [11]. Viruses were harvested and viral titrations were performed in Vero cells using plaque assay.

### Calu-3 cell infections and RNA isolation

All experiments involving infectious virus were performed in approved biosafety level 3 (BSL) laboratories at National Institute for Viral Disease Control and Prevention, China CDC. Cells were washed with MEM and inoculated with viruses at an multiplicity of infectious of infection (MOI) of 5 or mock-diluted in MEM for 2 h at 37°C. Following inoculation, cells were washed 3 times with MEM and fresh medium was added to signify zero hour. Triplicate samples of mock-infected and virus-infected Calu-3 cells were harvested at different times between 0 and 24 hour post-infection (hpi). Calu-3 cells were cultured for RNA isolation using Trizol reagent (Invitrogen, USA) following the manufacturer’s protocol.

### Library Construction and Sequencing

A total amount of 50 ng RNA per sample was used as input material for the Total RNA Library Construction and host rRNA removal according to the instructions of the Trio RNA-Seq kit (Nugen, 0506-32). Total RNA libraries were sequenced on the Illumina Novaseq using the 2×150bp paired-end read setting.

### Data analysis

Raw reads were filtered to obtain clean data by Trimmomatic (v0.35) (With parameters ‘ILLUMINACLIP:adapter.fa:2:30:10 HEADCROP:10 LEADING:3 TRAILING:3 SLIDINGWINDOW:4:15 MINLEN:36’)[12]. The cleaned data were mapped to the human GRCh38 reference genome using STAR aligner (v2.7.2a)[13]. The htseq-count command was used to count reads mapped to each gene [14]. The R package DESeq2 was applied to further identify Differentially Expressed Genes (DEGs) (FDR<0.05, |log2FC|>=1) [15]. The unmapped reads against the entire human genome were further aligned to the reference genome of SARS-CoV-2 (EPI_ISL_402119). Virus genome annotation was based on our previous work[16]. GO enrichment analysis was performed by Fisher’s exact test with the 19932 human protein-coding genes as a background in R. For analysis of microarray data of SARS-CoV and MERS-CoV, normalization and identification of DEGs (FDR<0.05, |log2FC|>=1) were conducted using the R package limma [17].

## Results

### Transcriptome profiling of virus-host interactions following SARS-CoV-2 infection

We carried out in time-course experiments to identify dynamic changes in transcripts in response to SARS-CoV-2 based on the infected and mock-infected groups across four time points (0, 7, 12 and 24 hpi), in which three biologically independent replicates for each treatment group were used for constructing cDNA libraries. The Calu-3 human airway epithelial cell line, a model of human respiratory disease [16], was used as the host cell of SARS-CoV-2, subjected to the same MOI and host cell used in the previous analyses of SARS-CoV and MERS-CoV infections. After the total RNA isolation and sequencing, we obtained the host transcriptomes, as well as the genomes and transcripts of viruses. The high-throughput sequencing resulted in an average of 49 million paired-end reads per sample, and the sequencing quality was high with a mean mapping rate of unique reads at approximately 72% among mock samples (Supplementary Table 1). The quality control of all samples was assessed by the PCA analysis based on normalized counts from DESeq2, which indicated that high quality was achieved given that the majority of samples were well clustered except only one sample from the infection group at 24 hpi that was removed before further analysis (Figure 1A).

**Figure 1.**
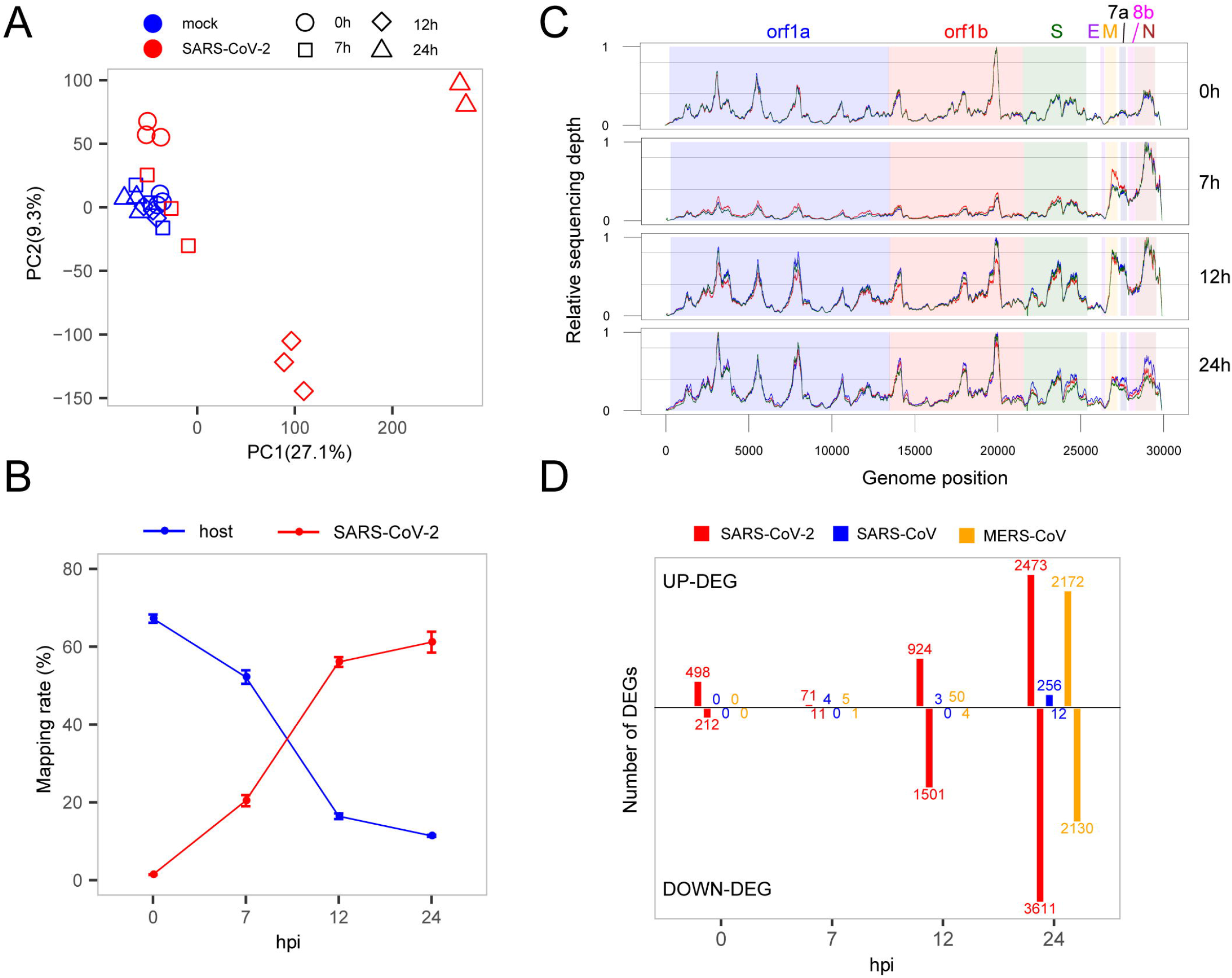
Interaction between SARS-CoV-2 and cell host. (A) PCA analysis of mock and SARS-CoV-2 infected samples. (B) Read mapping rate to the host or virus genomes. (C) Activity distribution of virus genome over times. The y-axis is relative sequencing depth that is normalized by (x-min)/(max-min) across the whole genome positions. Each line represents one biological replicate. (D) The numbers of DEGs at each time point among the three viruses. Only protein-coding genes were counted for SARS-CoV-2.

### Rapid growth of SARS-CoV-2 accompanied by dynamic changes of host genes

To evaluate the growth rate of SARS-CoV-2, we calculated the RNA level of the virus represented by unique reads mapping rates at different time points. Our results showed that in general the virus reads increased sharply from 1.4% to 61.2% while reads mapped to the host genome dropped rapidly from 67.2% to 11.4% (Figure 1B), suggesting a rapid replication of the virus within 24 hours. From the results, at the earliest time point (0 hpi), virus produced high-levels of viral genome RNA as evidenced by relatively even coverage depth across the whole genome (Figure 1C). Interestingly, we found that there was a significantly active transcription of the 3’ end of SARS-CoV-2 at 7 hpi, especially for the M, 6, 7a, 7b, 8b and N genes (Figure 1C) which could play important roles in the antagonism with host immune response[18, 19]. After that, the relatively even depth distribution of reads along viral genome was again observed at panels of 12 and 24 hpi. This time-dependent patterns of virus replication and transcription was most likely to play critical roles in the pathology of SARS-CoV-2.

To elucidate the global changes of host gene expression along with virus growth, we identified the overall up- and down-regulated DEGs during SARS-CoV-2 infection (Figure 1D and Supplementary Table 2). As shown in Figure 1D, during the early stage of infection before 7 hpi, there were many more up-regulated genes than down-regulated genes (498 vs 212 at 0 hpi, 71 vs 11 at 7 hpi), soon after, the number of down-regulated genes significantly exceeds that of up-regulated genes (924 vs 1501 at 12 hpi, 2473 vs 3611 at 24 hpi). Most importantly, most of DEGs at 0 hpi were suppressed at 7 hpi, which simultaneously occurred with active transcription of the 3’ end of SARS-CoV-2 genome, demonstrating the critical role of the 3’ end in antagonizing host immune response. The suppression of host responses were not likely due to sequencing bias because the three samples from the infected group at 7 hpi were clustered with mock samples (Figure 1A). Interestingly, there seem to be some correlated between the decrease in the levels of the host transcriptome (compared to the total RNA level of SARS-CoV-2) and the relative number of up-regulated genes (compared to down-regulated genes) (Supplementary Figure 1). This may indicate the complex molecular behavior of the host cell in response to the virus infection.

### Comparison of host transcriptome responses to SARS-CoV-2, SARS-CoV and MERS-CoV

To investigate specific host responses during SARS-CoV-2 infection, we performed a comparative transcriptome analysis by integrating two public host transcriptomes of SARS-CoV (GSE33267)[20] and MERS-CoV (GSE45042)[10] infected in the same cell line with the same MOI. Overall, a huge divergence was presented in time-specific DEG patterns among SARS-CoV-2, SARS-CoV and MERS-CoV (Figure 1D). For SARS-CoV-2, 710 DEGs (498 up-regulated and 212 down-regulated) were immediately induced at the very early stage (0 hpi), and many more DEGs were gradually observed at the late stages (12 and 24 hpi). In contrast, SARS-CoV and MERS-CoV infected cells exhibited far fewer DEGs (0, 4 and 3 for SARS-CoV and 0, 6 and 54 for MERS-CoV) at the early stages (0, 7 and 12 hpi). However, more DEGs were clearly detected at 24 hpi during SARS-CoV and especially MERS-CoV infection (268 and 4302 respectively). This distinct DEG patterns indicated that SARS-CoV-2 actually induced earlier host responses compared with SARS-CoV and MERS-CoV. To further delineate differential perturbation of pathways among three viruses, we conducted GO-enrichment analysis based on their respective DEGs. Overall, substantially enriched pathways, such as inflammation, apoptosis, antiviral response, transcription, translation and mitochondrion-related pathways, were detected at various time points during SARS-CoV-2 infection (Figure 2 and Supplementary Table 3). At 0 hpi, the up-regulated DEGs were mostly enriched in the pathways related to inflammation-related pathways including the NF-kB signaling and cytokine-mediated signaling pathways, suggesting that SARS-CoV-2 could induce inflammatory responses at the very early stage of infection. At the same time, SARS-CoV-2 also triggered the cellular apoptosis signaling pathway, implying that early onset cell death happened along with inflammation response. Beginning at 7 hpi, our results showed a significant enrichment in antivirus response-related pathways until 24 hpi (Figure 2). At the late stages (12 and 24 hpi), down-regulated DEGs were exclusively enriched in fundamental host pathways responsible for RNA processing and transcription, protein translation and mitochondrial activity (Figure 2). Different from SARS-CoV-2, at the late stage of SARS-CoV infection (24 hpi), the highly enriched genes were identified to be involved in antivirus-related pathways, whereas no significantly enriched pathways were found for MERS-CoV infection despite numerous DEGs existing at 24 hpi (Figure 2). Taken together, the above results indicated that the etiology mechanism of SARS-CoV-2 was different from that of SARS-CoV and MERS-CoV as implicated by the overall differential patterns of the host response against infection.

**Figure 2.**
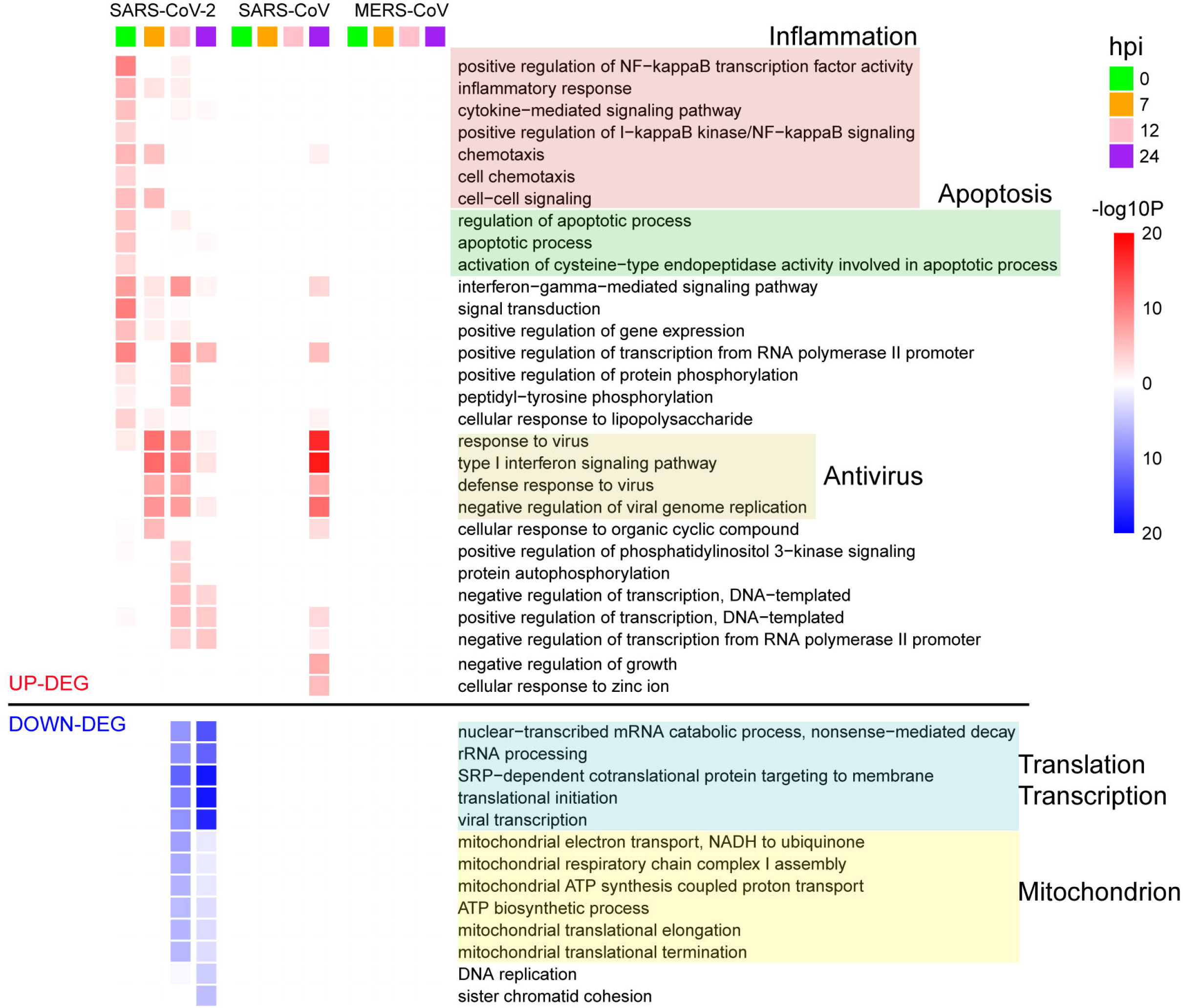
GO enrichment analysis of DEGs for the three viruses. The GO BP terms with enrichment FDR<0.001 are shown.

### Quantification of the capacity for host antiviral immunity and cytokine production for SARS-CoV-2, SARS-CoV and MERS-CoV infections

As mentioned above, SRAR-CoV-2 induced specific patterns of host antiviral and inflammation responses compared to SARS-CoV and MERS-CoV. To quantify host antiviral capacity and inflammation responses during infection of the three viruses, two sets of genes were used as their indicators. First, we used a set of 45 early induced genes in interferon-α treated Calu-3 cell [7] as antiviral indicators to quantify the level of host antiviral capacity against SARS-CoV-2, SARS-CoV and MERS-CoV infections. Our analysis showed that, the antiviral capacity of the host against SARS-CoV-2 was gradually increased over the time course of infection (Figure 3A). In contrast, the host antiviral capacities against both SARS-CoV and MERS-CoV were nearly zero at least during the initial stages of infection (between 0h and 12 hpi), followed by a marginal increase at 24 hpi. The antiviral capacity in SARS-CoV and especially MERS-CoV infected cells were much lower than that in SARS-CoV-2 infected cells, which might underpin the disparity in mortality between the three viruses. Despite the observation of the potent early-induced host antiviral activity during SARS-CoV-2 infection as compared to SARS-CoV and MERS-CoV infection, our results clearly showed that most of the genes (25/45) were significantly induced among infections of the three viruses (Figure 3B). In addition, a list of the virus-specific antiviral-related genes was identified, including PARP10[21] and CMPK2[22] for SARS-CoV-2, BST2[23], ITITM1 and USP41 for SARS, and PARP4[24] for MERS-CoV (Figure 3B).

**Figure 3.**
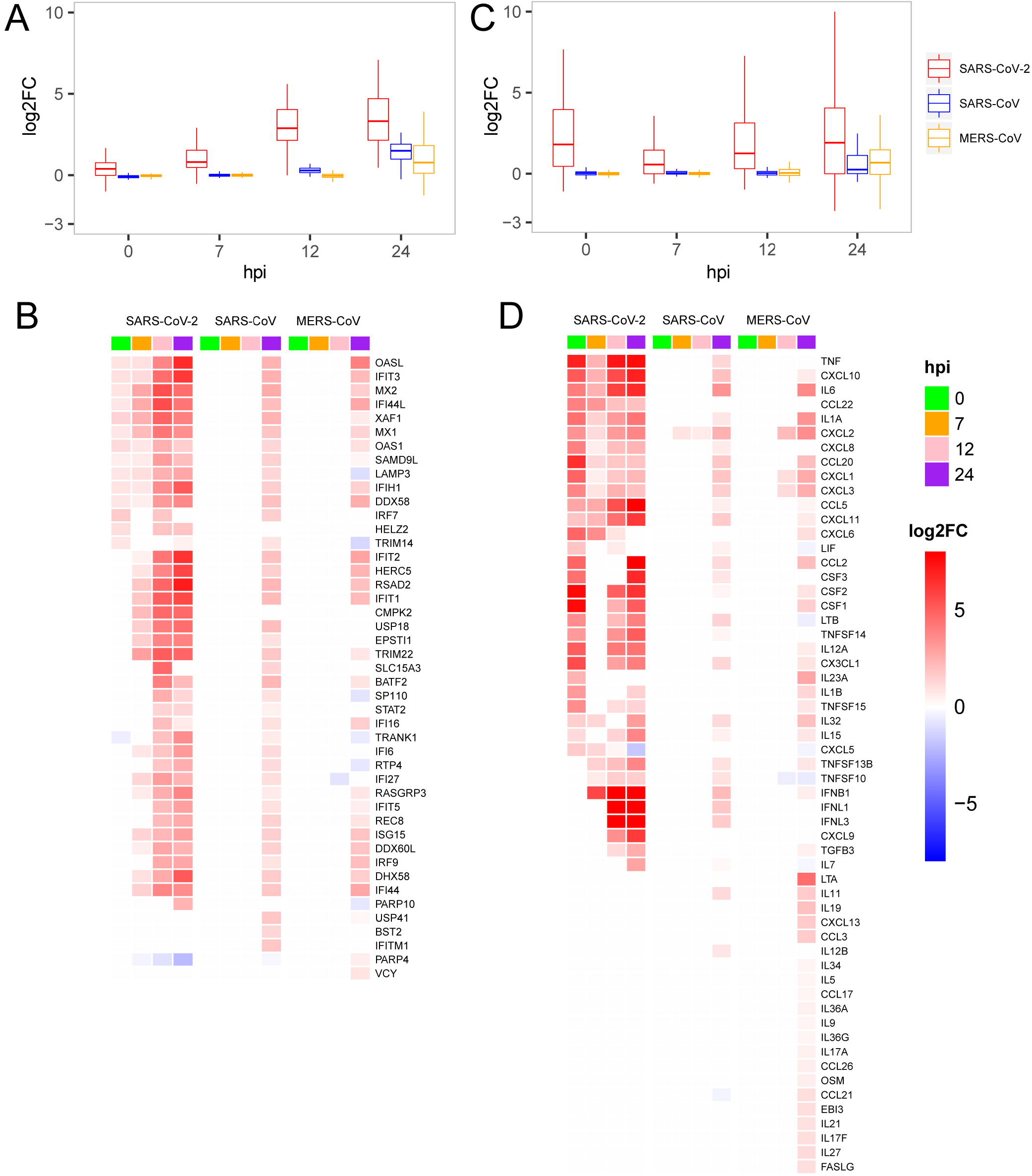
Expression patterns of the host antiviral-related genes and cytokines. (A) Quantification of host antiviral capacity. (B)Expression patterns of the host antiviral-related genes. (C) Quantification of the host cytokine genes. (D) Expression patterns of the host cytokine genes.

Secondly, we further used a set of 113 human cytokines to quantify host inflammation responses between three viruses. The 113 cytokines from the CytoReg database were often cited by various publications and play a primary role in the immune system[25]. Our results showed that, for SARS-CoV-2, the level of cytokine production was highly induced at 0 hpi, decreased at 7 hpi, and then slowly recovered thereafter (Figure 3C). Relatively high levels of cytokine expression only occurred at 24 hpi for SARS-CoV and MERS-CoV. Our analysis also provided evidence that SARS-CoV-2 had more cytokines in common with SARS-CoV than with MERS-CoV (Figure 3D). Unlike the other two viruses, MERS-CoV specifically induced the expression of dozens of cytokines, such as LTA, IL19, CXCL13 and CCL3, at 24 hpi, which were not observed in the case of the other two viruses. Interestingly, among the 28 up-regulated cytokines at the very early stage (0 hpi) during SARS-CoV-2 infection, eight cytokines including IL-6 (IL6), IL-1b (IL1B), IL-8 (CXCL8), G-CSF (CSF3), GM-CSF (CSF2), IP10 (CXCL10), MCP1 (CCL2) and TNF were reported to exhibit substantially elevated serum levels [9, 26, 27], which indicated that early induction of cytokines played critical roles in the pathology of SARS-CoV-2. While most of the eight cytokines were moderately up-regulated at the late stage during SARS-CoV and MERS-CoV infections, up-regulation were not observed at the early stage. Collectively, SARS-CoV-2 induced distinct patterns of host antiviral response and cytokine production.

### Regulation of key genes from cell entry to type-Ⅰ interferon production

Next, to gain possible explanations for the distinct patterns in host antiviral capacity and cytokine production during SARS-CoV-2 infection, dynamic expression of four types of key genes were evaluated, including virus receptors for cell entry, pathogen recognition receptors (PRRs) for an innate immune startup, regulator genes for induction of antiviral-related genes and interferon production (Figure 4). For the three cell entry related genes (ACE2 as the receptor of SARS-CoV and SARS-CoV-2 [28, 29], DPP4 as the receptor of MERS-CoV [30] and protease TMPRSS2 for S protein priming of SARS-CoV-2 [27]), we observed the dramatic changes in TMPRSS2 expression with very early induction during SARS-CoV-2 infection, and the slightly down-regulated expression of ACE2 in cells infected with SARS-CoV-2 and SARS-CoV, whereas DPP4 was more up-regulated in MERS-CoV (Figure 4). For the two PRRs, DDX58 is a canonical RIG-I-like receptor for RNA virus recognition[31], and TLR3 is a Toll-like receptor playing important roles in initiating a protective innate immune response to highly pathogenic coronavirus infections[32]. We observed that all three viruses had a notably up-regulated expression of DDX58 while only MERS-CoV had a suppressed TLR3 at the 24 hpi (Figure 4), which is consistent with the fact that decreased expression of TLR3 contributes to the pathology of highly pathogenic coronavirus infections[32]. Among the four regulator genes, IRF7 is responsible for the expression of most IFN-α subtypes and the type I IFN amplification loop[33], and IRF9, STAT1 and STAT2 form the ISGF3 complex that binds to interferon-stimulated response elements and thereby induces the expression of interferon-stimulated genes[34]. As expected, gradually up-regulation of the four primary regulator genes was observed for all three viruses (Figure 4). At last, we found a significant difference in the expression of IFNB1 between SARS-CoV-2, SARS-CoV and MERS-CoV, indicating that IFNB1 likely accounted for the observed variations of the host antiviral capacities among three viruses (Figure 4). Taken together, early induction of TMPRSS2 and gradually increased expression level of IFNB1 were likely responsible for the distinct host immune response patterns of SARS-CoV-2 infection.

**Figure 4.**
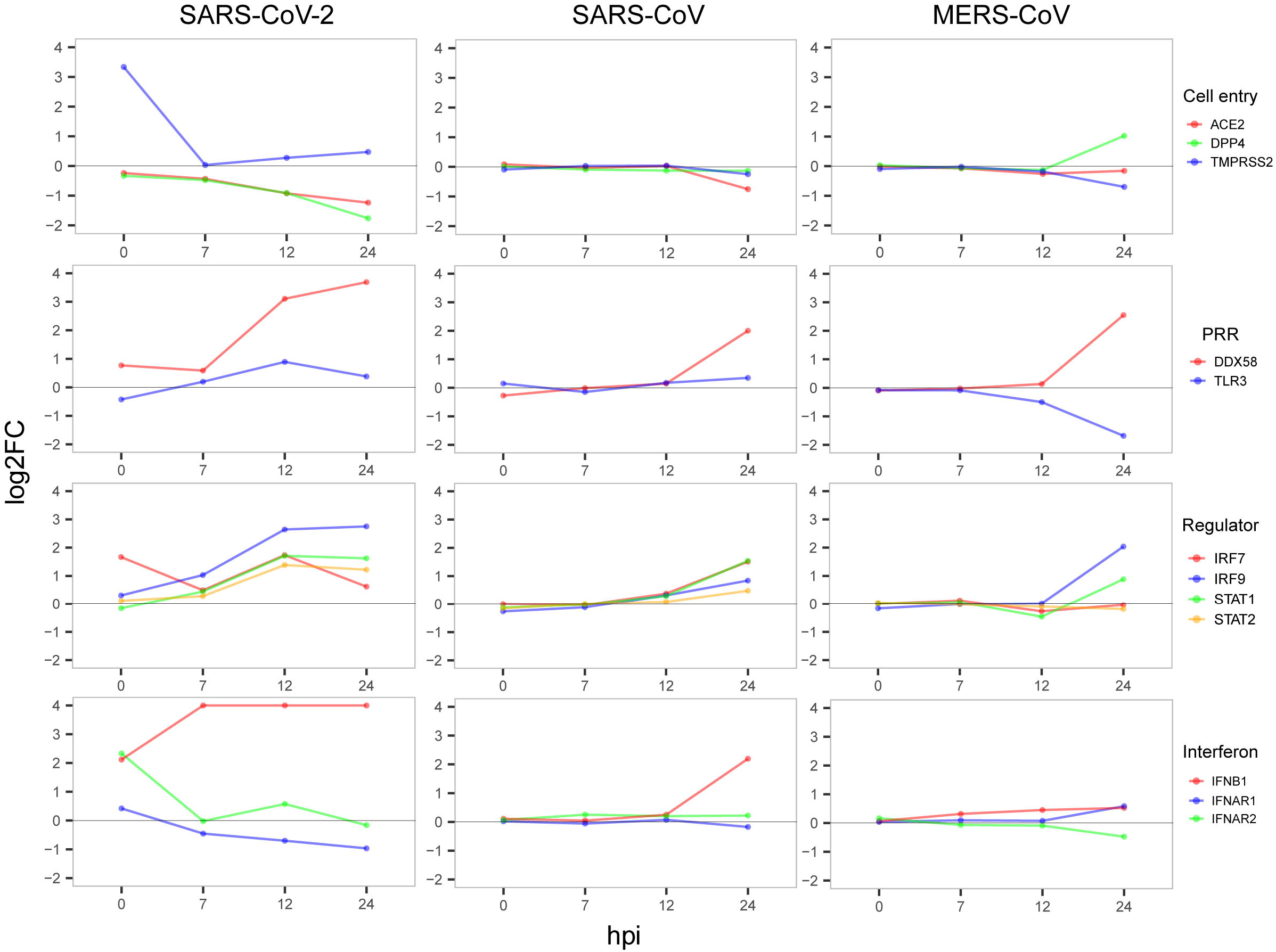
Dynamic expression of four types of important genes. The value of y-axis was restricted to have a maximum of 4 to show notable gene expression changes.

## Conclusion

Using time-series profiling of the virus genome and host transcriptome at the same time during SARS-CoV-2 infection coupled with comparative transcriptome analysis, we found that, compared to SARS-CoV and MERS-CoV, SARS-CoV-2 induces strong host cell responses at the very early stage of infection that not only favor its high infectivity to host cells but also restrict its pathogenesis.

## Discussion

Here we sequenced the transcriptomes of SARS-CoV-2 and virus-infected host cells simultaneously during the early stages of infection, providing a robust reference dataset to speculate the antagonistic pattern between pathogen and host cells. To summarize, our findings showed that SARS-CoV-2 induced the significantly high expression of the cellular serine protease TMPRSS2 at 0 hpi to help the entry of viral particles into cells[28] (Figure 4). At the same time, host cell initiated an immediate response for the invasion of SARS-CoV-2 virus (Figure 1D). Then, the virus successfully suppressed the acute response of host cells for fast proliferation by increasing the transcripts of its 3’ genome end, including M, 6, 7a, 7b, 8 and N genes which were consistent with their reported regulations to host immune response[18, 19]. As a response from hosts cell, a number of antiviral pathways and cytokine productions were up-regulated to resist the virus infection (Figures 2 and 3). In particular, several metabolism-associated pathways were down-regulated at 12hpi and 24 hpi (Figure 2). After the antagonistic cycle, a dramatic proliferation of viral particles was detected in the early infection of host cells (Figure 1B), which could possibly be an explanation for the fast spread of SARS-CoV-2 in humans.

As SARS-CoV-2 was reported a relatively low risk of mortality[3] compared to the other two serious human coronaviruses, SARS-CoV and MERS-CoV, we compared and contrasted the host transcriptomes in response to the viral infections. We found that some cytokines in SARS-CoV-2-infected cells were markedly up-regulated at a very early stage, which was not observed for SARS-CoV and MERS-CoV and even less frequently observed for other viruses. The unusual high expression of cytokines at 0 hpi possibly explains why patients with severe clinical symptoms rapidly deteriorated. Although the number of infected cases was very high, the majority of infections displayed mild symptoms which are partly explained by a gradual increase in host antiviral capability from 7 to 24 hpi. In contrast to SARS-CoV-2, both SARS-CoV and MERS-CoV were able to inhibit the antiviral capability of the host significantly, which could explain their observed relatively high mortalities. MERS was associated with a higher mortality than SARS, which could be in part attributed to the higher expression of cytokines suppressing the antiviral responses.

Recently, Blanco-Melo et al. [5] have published transcriptome data of host responses to SARS-CoV-2 from *in vitro* cell lines including A549 (MOI of 0.2) and NHBE (MOI of 2) at 24 hpi. This previously published data is complemented by our study designed to investigate the early response phase of cell lines infected with SARS-CoV-2. While the previous work did not observe the elevated levels of IFNB1, IFNL1 and IFNL3, our findings show that not only IFNB1 but also IFNL1 and IFNL3 expressions are up-regulated between 7 and 24 hpi (Figure 3D). Also, they did not detect gene expression of ACE2 and TMPRSS2 at 24 hpi, while we observed that ACE2 is down-regulated at 24 hpi and TMPRSS2 is only up-regulated at 0 hpi before returning to the normal levels (Figure 4). Our time-series sampling revealed distinct early-response features of SARS-CoV-2, which provided a possible explanation for some clinical observations. For example, a recent clinical study [35] found that SARS-CoV-2 could replicate effectively in upper respiratory tract tissues, and that the viral loads appeared earlier (before day 5) and were substantially more than expected. Findings from the present study have confirmed that, at 7 hpi, the 3’ end of SARS-CoV-2 genome start to express densely, reducing the effectiveness of host immune surveillance, which possibly enables the rapid replication of SARS-CoV-2 in upper respiratory tract tissues.

In spite of the fact that several studies have already demonstrated a consistent correlation between gene expression measured by RNA-Seq and by microarray [36–38], we still need to exclude the possibility of bias resulting from different methodologies. First, because RNA-Seq can potentially detect more genes than microarrays, we only considered protein-coding genes for the analysis of RNA-Seq results. For SARS-CoV-2, the microarray analysis identified more than 90% of the 6800 DEGs, including 6514 DEGs of SARS-CoV and 6198 DEGs of MERS-CoV. Secondly, expressions of the 6800 DEGs were distributed over the four time points from low to high, not only in SARS-CoV-2 but also in SARS-CoV and MERS-CoV (Supplementary Figures 2 and 3), indicating that the silent early host responses to SARS-CoV and MERS-CoV appeared not to be due to technological biases. Lastly, when extending the infection time from 24 hpi to 72hpi (GSE33267), thousands of DEGs (minimum 1022 and maximum 2017 genes), which had been inhibited at the early stages, were actually induced (Supplementary Figure 4).

## Acknowledgments

This work has been supported by the National Key Research and Development Program of China (2016YFD0500301) and the CAMS Initiative for Innovative Medicine (grant 2016-I2M-1-005), and by the National Natural Science Foundation of China (grants 31671371 and 31601082), and by the Central Public-Interest Scientific Institution Basal Research Fund (2017PT31026, 2018PT31016).

## Data availability

The sequencing data from this study have been submitted to the National Genomics Data Center (https://bigd.big.ac.cn/) with the accession number PRJCA002617.

## Contributions

Jiang, Tan and Huang devised the experiment and wrote the paper; Sun, Wu, Sheng and Zhu conducted bioinformatics analysis; Ye, Huang and Yang prepared samples; Wang provided Calu-3 cell; Pan run RNA Sequencing; and all authors revised the manuscript.

---

**Supplementary Figure 1.** Variation of the number of up-regulated genes minus down-regulated genes over time.

**Supplementary Figure 2.** Expression levels of cytokine genes between mock and infected groups. The left column is for SARS-CoV-2, the middle column for SARS-CoV and the right column for MERS-CoV. For SARS-CoV-2, the expression level is quantified by log2(TPM+1). Cytokine genes are highlighted.

**Supplementary Figure 3.** Expression levels of antivirus-related genes between mock and infected groups. The left, middle and right columns show results for SARS-CoV-2, SARS-CoV and MERS-CoV respectively. For SARS-CoV-2, the expression level is quantified by log2(TPM+1). Antiviral-related genes are highlighted.

**Supplementary Figure 4.** Variation of the number of DEGs during SARS-CoV infection.

