## Supplementary figures and images for "Comparative transcriptome analysis reveals the intensive early-stage responses of host cells to SARS-CoV-2 infection"

### Supplementary Figure 1

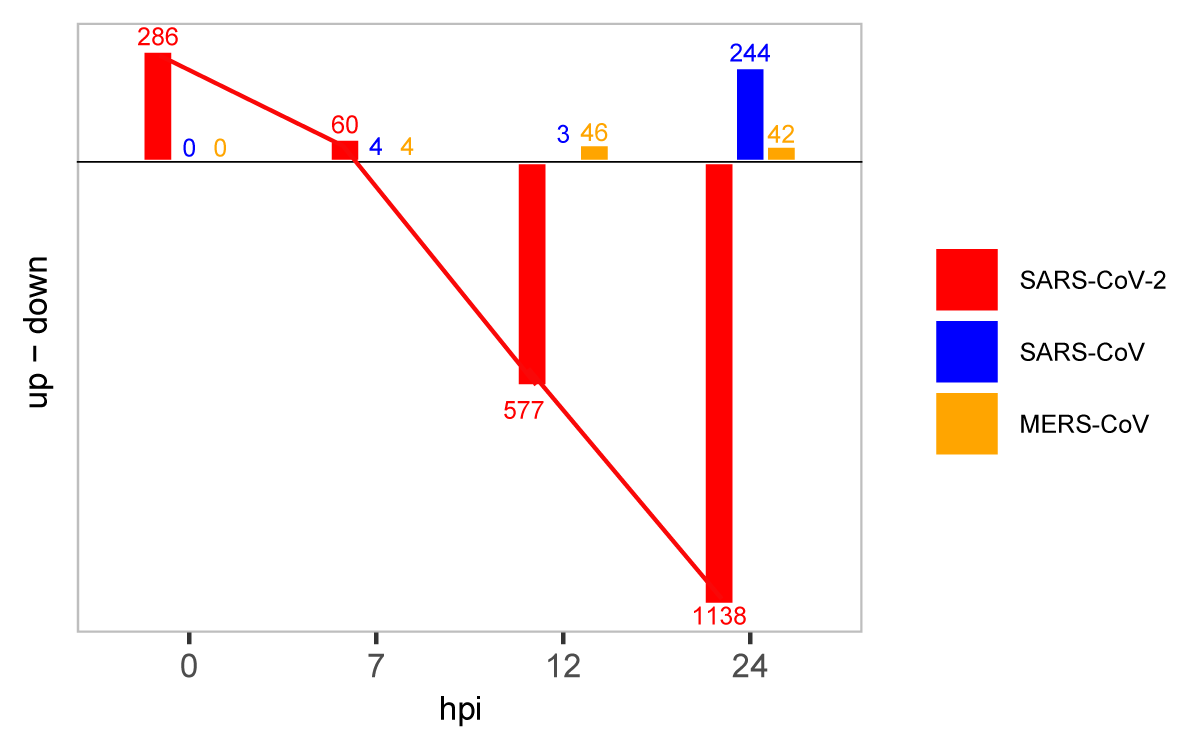

### Supplementary Figure 2

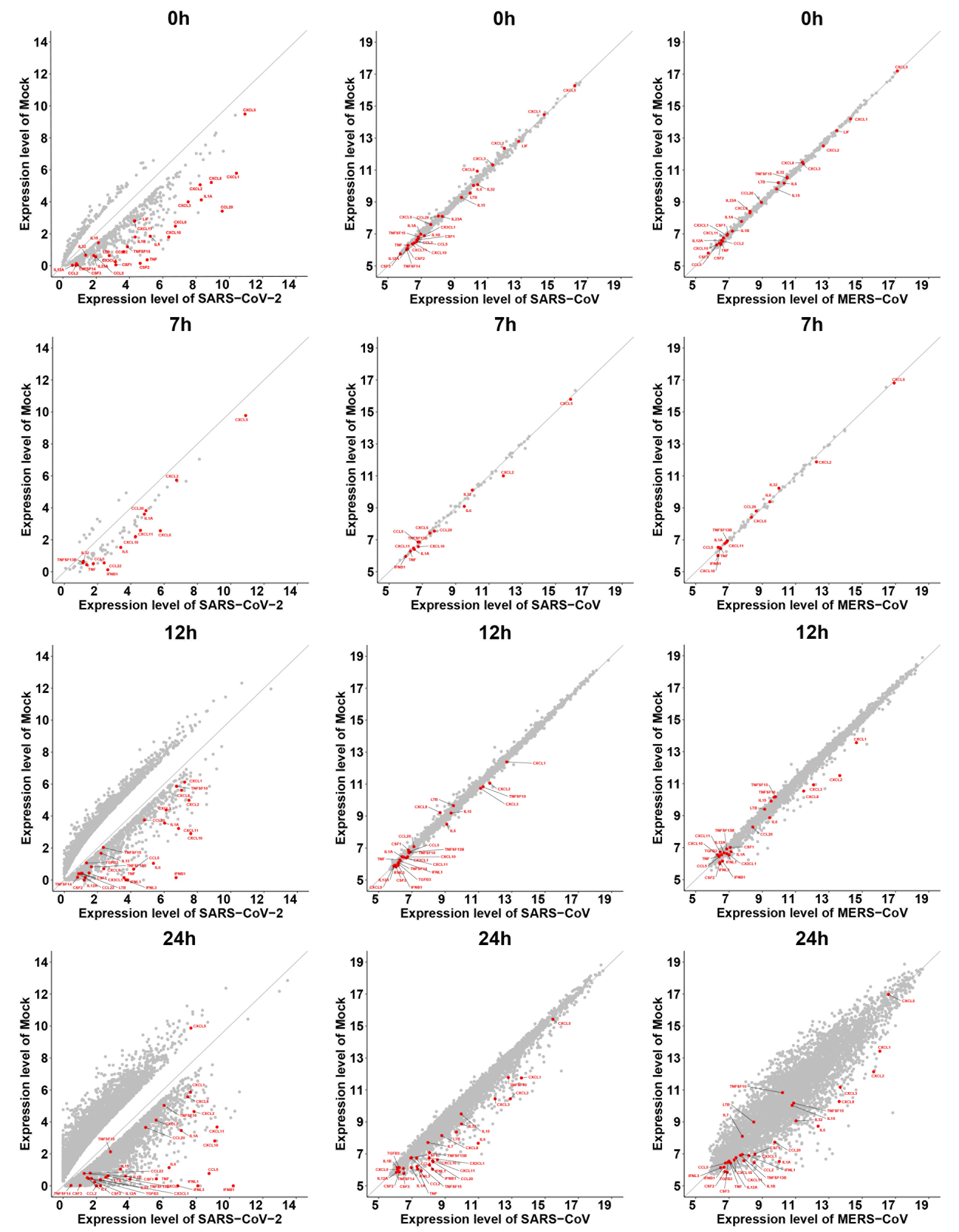

### Supplementary Figure 3

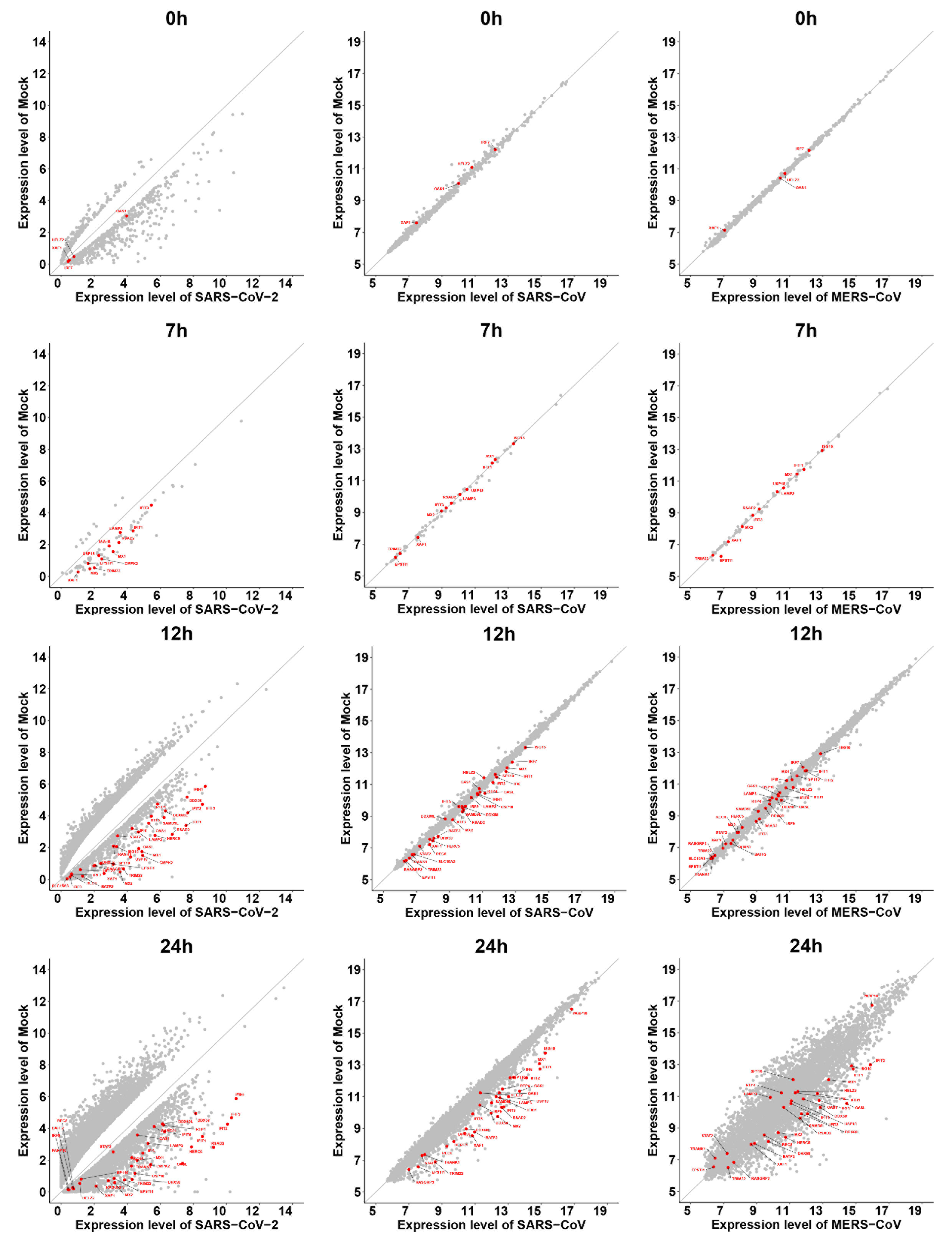

### Supplementary Figure 4

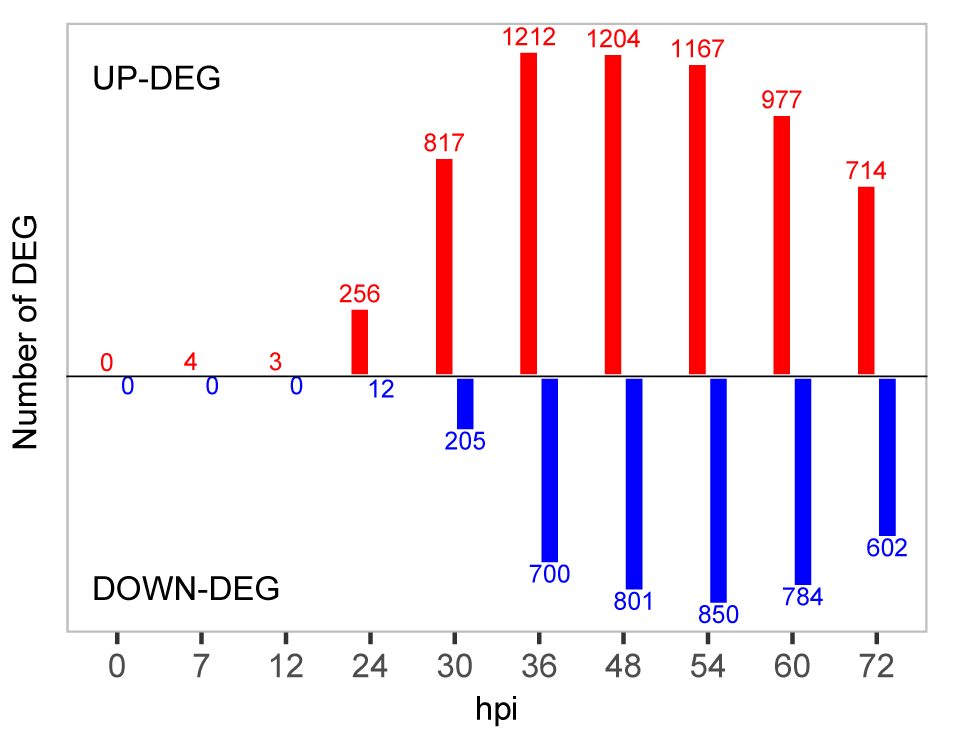
